# Deriving Ranges of Optimal Estimated Transcript Expression Due to Non-identifiability

**DOI:** 10.1101/2019.12.13.875625

**Authors:** Hongyu Zheng, Cong Ma, Carl Kingsford

## Abstract

Current expression quantification methods suffer from a fundamental but under-characterized type of error: the most likely estimates for transcript abundances are not unique. This means multiple estimates of transcript abundances generate the observed RNA-seq reads with equal likelihood, and the underlying true expression cannot be determined. This problem is called non-identifiability for probabilistic models, and is further exacerbated by incomplete reference transcriptome. That is, reads may be sequenced from unannotated expressed transcripts. Graph quantification is a generalization to transcript quantification, accounting for the reference incompleteness by allowing exponentially many unannotated transcripts to express reads. We propose methods to calculate a “confidence range of expression” for each transcript, representing its possible abundance across equally optimal estimates for both quantification models. This range informs both whether a transcript has potential estimation error due to non-identifiability and the extent of the error. Applying our methods to the Human Body Map data, we observe 35%–50% of transcripts potentially suffer from inaccurate quantification caused by non-identifiability. When comparing the expression between isoforms in one sample, we find that the degree of inaccuracy of 20%–47% transcripts can be so large that the ranking of expression between the transcript and its sibling isoforms cannot be determined. When comparing the expression of a transcript between two groups of RNA-seq samples in differential expression analysis, we observe that the majority of detected differentially expressed transcripts are reliable with a few exceptions after considering the ranges of the optimal expression estimates. The code for computing the range of expression is available at https://github.com/Kingsford-Group/subgraphquant. The code for the involved analyses is available at https://github.com/Kingsford-Group/subgraphquantanalysis.

## 1 Introduction

Despite the improvements of transcript expression estimation methods based on RNA-seq data [1–4], the estimated transcript expression can still be inaccurate and uncertain. One source of uncertainty in expression estimation is that multiple sets of expression estimates can optimally explain the observed RNA-seq reads. Therefore, the “best” estimation cannot be uniquely identified. The state-of-the-art methods to quantify transcripts’ expression are based on probabilistic models, and, in probabilistic model inference terminology, the phenomenon of non-uniqueness in optimal parameters under infinite data is called model “non-identifiability”. In this work, we relax the concept and use this term to refer to the non-uniqueness of optimal parameters under a given finite dataset. See Figure 1 for a toy example of model non-identifiability in expression quantification. Two main problems for evaluating the accuracy of transcript expression estimates under non-identifiability are (1) detecting the transcripts whose expression estimates are unidentifiable and (2) bounding the range of the uncertain expression of the transcripts.

**Figure 1:**
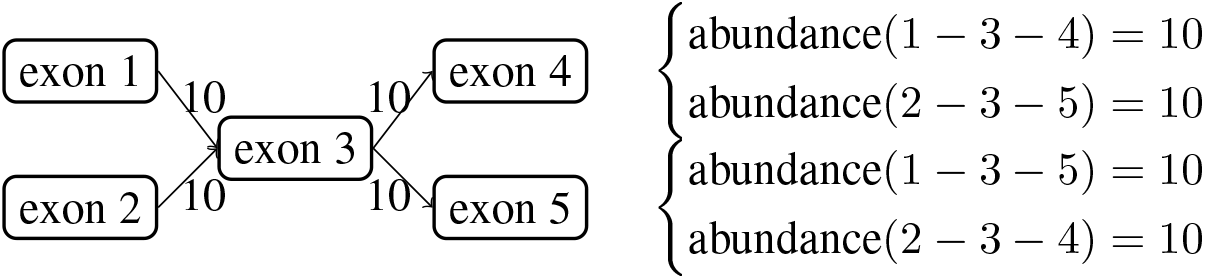
Example of a non-identifiable quantification model with four transcripts. Transcripts are the paths in the splice graph, denoted by the concatenation of exon indices. The number on each edge indicates the number of observed reads mapped to this splice junction. The set of transcript abundances are optimal if it perfectly explains the observed reads. That is, for each junction, the total abundances of transcripts containing that junction sum up to the number displayed on the edge. The right side of the figure shows two co-optimal expression abundances. It can be verified that both solutions explain the observed reads perfectly as they both predict 10 reads on each junction.

Expression of transcripts is used in many analyses, and understanding the accuracy and uncertainty of the estimated expression helps us evaluate the confidence of the conclusions of such analyses. Transcript expression estimates are used for detecting splicing isoform switching [5–7], for identifying differential expression [8–12], and for predicting disease status and treatment outcome [13, 14]. Ranges of uncertain expression estimates provide useful insights into the the reliability of these studies.

We focus on the uncertainty of expression estimates due to model non-identifiability, but there are other causes of estimation uncertainty or inaccuracy. The small sample sizes (low sequencing depth of RNA-seq) also leads to estimation errors in transcripts’ expression. Statistical methods such as bootstrapping and Gibbs sampling [15–17] provide an estimate on the error in expression levels due to the sample size. This estimation error can be reduced by increasing the sequencing depth. In contrast, the estimation uncertainty due to model non-identifiability is more fundamental because it cannot be addressed even under infinite sample size. In RNA-seq data, exon sequences are usually shared among multiple transcripts, and RNA-seq reads usually cannot be mapped to a unique transcript. Due to the high occurrence rate of multi-mapping events in RNA-seq data, the best set of transcripts’ expression cannot be uniquely resolved, and the phenomenon of non-identifiability occurs. Multi-mapped reads are prevalent in RNA-seq data regardless of the sequencing depth. Thus the uncertainty of expression estimated due to model non-identifiability cannot be easily addressed.

Previous works have analyzed the non-identifiability phenomenon in transcript expression quantification. However, they mainly focused on the first problem, to identify the transcripts for which the expression estimates are unidentifiable, but not the second, which is to bound the ranges of the true expressions for the transcripts with uncertain expression estimates. Lacroix et al. [18] and Hiller et al. [19] developed methods to list the transcripts that have non-unique expression estimates but their methods do not provide information about the values of optimal abundances. Roberts et al. [20] incorporated the detection of non-identifiable abundances into their quantification method, and designed a tool to output an identifiability flag for each transcript along with a single expression estimate.

In this work, we develop methods that address the range of optimal expression estimates for transcripts under model non-identifiability. For each transcript, we calculate the minimum and maximum values across all optimal expression estimates. That is, for any expression value between the computed minimum and maximum, there exists a set of expression of the other transcripts such that the estimation of all transcripts’ expression (combining both the expression of this transcript and the other transcripts) leads to the largest likelihood in the probabilistic model of expression quantification. Compared with a list of names of the transcripts for which the expressions are unidentifiable, the range of optimal expression of a transcript provides more information on the accuracy (or inaccuracy) of the estimate.

Most widely used quantification software [1–4] take a set of reference transcript sequences as input and assume the reference transcripts are the complete set of sequences that can be expressed. This is called reference-transcript-based expression quantification. Another line of expression quantification models [21– 23] called “graph quantification” assumes that the current reference transcriptome database is incomplete. Instead, the splice graph which encodes the exon-exon connection relationships is assumed to be correct and complete, meaning every possibly expressed transcript corresponds to a path in the splice graph and vice versa. Those models infer the splice graph edge expression or edge selection propensity in the quantification probabilistic models. We provide a more detailed overview in Section 2.3. Many transcript assembly methods also adopt a similar setup, assuming a mixture of reference transcripts and novel isoforms are expressed [24–28].

We develop methods to bound the range of optimal expression estimates for both reference-transcript-based and graph quantification models. Our method for the reference-transcript-based quantification is based on linear programming over sufficient statistics (Section 2.2), and for graph quantification is based on max-flow formulations to “introspect” the graph quantification model (Section 2.4). Our introspection algorithm can not only bound the uncertain expression of full transcripts, but also extends to graph structures. For example, given a set of edges, our method computes the range of the optimal total expression of transcripts that cover any edge in the set.

Combining our methods for quantification models and interpolating between the complete and incomplete reference transcriptome assumption, we can additionally compute the range of optimal expression estimates under the assumption that a given percentage of the expression comes from the reference and the remaining expression come from the full paths in splice graphs (Section 2.5).

Applying our method to 16 Human Body Map samples, we analyze to what degree the expressions of transcripts are estimated inaccurately due to non-identifiability. We observe that around 35%–50% of transcripts potentially suffer from expression estimation error across the 16 samples. Most of these transcripts (or 20%–47% of total transcripts) have very uncertain expression estimates such that the ranking of expression between the transcript and its sibling isoforms from the same gene is inconclusive. Around half of the transcripts with uncertain expression estimates due to non-identifiability are different from those due to finite sample size.

Applying our method on sequencing datasets of a MCF10 cell line and of CD8 T cells, we use the ranges of optima to evaluate the reliability of detected differentially expressed transcripts within each dataset. A DE detection is unreliable if the ranges of optimal expression between DE groups largely overlap. We observe that the majority of the DE calls are reliable and robust to the uncertain expression estimation due to non-identifiability when the reference transcriptome contributes to more than 40% of expression. But there are 5 unreliable DE calls (out of 257 detections) in the MCF10 dataset and 19 unreliable DE calls (out of 3152 detections) in the CD8 T cell dataset. It requires further investigation to determine whether these transcripts are actually differentially expressed, and analyses based on the DE status of these transcripts require extra caution.

## 2 Methods

We start with relevant definitions in 2.1. Section 2.2 provides a high-level overview of probabilistic modeling of transcript quantification, and the linear programming to derive range of optimal abundance estimates for this setup. Section 2.3 provides an overview of graph quantification analogous to previous section, and Section 2.4 describes our introspection algorithm to derive the ranges under this setup.

### 2.1 Definitions

A “splice graph” is a directed acyclic graph representing alternative splicing events in a gene. The graph has two special vertices: *S* represents start of transcripts and *T* represents termination of transcripts. Every other vertex represent a (partial) exon. Edges in the splice graph represent “splice junctions,” i.e. potential adjacency between exons in transcripts. Each transcript corresponds to a unique *S* − *T* path in the splice graph, and the set of known transcripts is called the “reference transcriptome”. *S* − *T* paths that are not present in the reference transcriptome correspond to “unannotated transcripts”. We use the phrase “quantified transcript set” to denote a set of transcripts with corresponding abundances.

### 2.2 Ranges of Optimal Estimates for Reference-Transcript-Based Quantification

Recall the problem of reference-transcript-based expression quantification. We focus on paired-end reads for now, however this formulation naturally extends to other types. We also focus on a particular probabilistic modeling, which lies at the heart of most modern transcript quantifiers [1–4]. Assume the paired-end reads from an RNA-seq experiment are error-free and uniquely aligned to a reference genome as fragments. We denote the set of fragments as *F*, the set of transcripts as 𝒯= {*T*_1_, *T*_2_, …, *T*_*n*_} with abundance (copies of molecules) *c*_1_, *c*_2_, …, *c*_*n*_. The probability of observing *F* is:

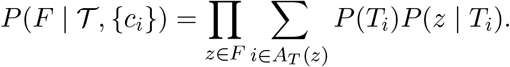

To generate a fragment, first a transcript is sampled, then the fragment is sampled from the selected transcript, and we only observe the resulting fragments. *P* (*T*_*i*_) denotes the probability of sequencing a fragment from transcript *T*_*i*_, and *P* (*z* | *T*_*i*_) denotes the probability of sampling the fragment *z* given that it comes from *T*_*i*_. *A*_*T*_ (*z*) is the set of transcript indices onto which *z* can map. Usually, *P* (*T*_*i*_) ∝ *c*_*i*_, and *P* (*z* | *T*_*i*_) is a known quantity derived from fragment length distribution, bias correction and similar factors. Recent transcript quantifiers [3, 4] employ complicated techniques to ensure accurate estimation of *P* (*z* | *T*_*i*_) and fast inference of {*c*_*i*_}.

We now introduce the idea of reparameterization. Assume, for now, each fragment spans exactly one junction. If two sets of quantified transcripts result in exactly the same total abundance on each junction, they yield the same generative model. In other words, these two sets will be indistinguishable from each other. Let {*g*_*i*_} be the set of total abundances for each junction *J*_*i*_, and we use *J*_*i*_ *∈ T*_*j*_ to denote that junction *J*_*i*_ is in transcript *T*_*j*_. The following condition is sufficient to guarantee that the set of transcript abundances 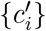 is valid, and indistinguishable from {*c*_*i*_}:

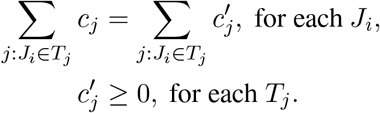

As this defines a linear system, we calculate the maximum and minimum possible value of 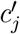 for each transcript *T*_*j*_, which would be the confidence interval for this transcript assuming {*c*_*i*_} is an optimal solution for the inference, which can be readily obtained using existing software. More specifically, we use either max 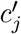 or min 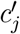 as the sole optimization objective with the linear system as the constraints, then solve the resulting linear program, for each transcript of interest. It is an interval because for every abundance value in between the extremes, there exists an optimal solution that allocates this exact abundance to the transcript.

This argument no longer holds when fragments can span multiple junctions. However, as we have shown previously [23], to ensure that two sets of quantified transcripts yield the same fragment distribution, it suffices that for each phasing path (that is the set of junctions in a fragment, as a path on the splice graph) the total abundance of transcripts containing the phasing path in full are equal. In other words, the form of the linear program remains unchanged, with the only change being that *J*_*i*_ is now a phasing path instead of a single junction.

### 2.3 Bird’s-Eye View: Graph Quantification

We now provide a high-level overview of graph quantification, and draw similarities with its transcript-based counterpart. The problem of splice graph expression quantification is very similar to the transcript version of the problem, but with the variable associated with transcripts {*T*_*i*_}, {*c*_*i*_} replaced by a graph *G* = (*V*, {*e*_*i*_}) and a network flow {*f*_*i*_} on the graph. We start by assuming that *G* is the splice graph in the usual sense, and every fragment *z* is a junction read, which can be mapped to an edge in *G*. Under an analogous probabilistic model, the probability of observing *F* is:

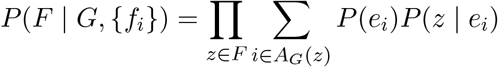

To generate a fragment, first an edge in *G* is sampled, then the fragment is sampled from the selected edge and we only observe the resulting fragments. Again, *P* (*e*_*i*_) denotes the probability of sampling a fragment from edge *e*_*i*_, and *P* (*z* | *e*_*i*_) denotes the probability of sampling the fragment *z* given it comes from *e*_*i*_. *A*_*G*_(*z*) denote the set of edges *z* can be generated from. We also have *P* (*e*_*i*_) ∝ *f*_*i*_ up to a normalization factor, and *P* (*z* | *e*_*i*_) being a known quantity.

After inferencing the splice flow value from the set of observed fragments, we obtain the optimal splice graph flow that explains the observation. While the graph flow itself finds use in certain cases, for most analysis it is imperative that the flow will be decomposed into weighted set of *S* − *T* paths, corresponding to a quantified set of transcripts. Importantly, the flow decomposition is not unique in many cases, and while these decompositions lead to different set of transcripts, they define the same generative model. This is a direct manifestation of non-identifiability, if we consider the transcript abundances to be the variables of the probabilistic model. In the next section, we will describe methods that consider all possible flow decompositions of a given graph flow. As a particular use case, these methods are able to determine the minimal and maximal possible abundance for a given transcript, across all possible flow decompositions.

Similarly to the transcript-based quantification, the conclusion no longer holds when fragments span multiple junctions, and it is much harder to adjust the graph quantification model to properly take these into account. However, for graph quantification *G* can be any supergraph of the splice graph, as long as each *S* − *T* path in *G* corresponds to a valid transcript to generate fragments, and each valid transcript corresponds to a unique *S* − *T* path in *G*. Several approaches are possible, and we will use the one outlined in our previous work [23], which properly handles phasing reads.

### 2.4 Subgraph Quantification

As discussed in the last section, non-identifiability manifests itself in graph quantification as different flow decompositions from the inferred splice graph flow. Specifically for a transcript, the corresponding *S* − *T* path can be assigned different weights from different flow decompositions. Similar to Section 2.2, we only need to know the minimal and maximal possible weight to construct the confidence interval.

Additionally, we are also interested in the related problem of “local quantification”, that is to estimate the abundance of a specific splicing pattern by aggregating expression of full-length transcripts containing the pattern. This is natural for analysis of complex gene loci, where certain splicing patterns within a region might be of larger interest than the patterns outside. Our reasoning for deriving a “confidence interval” can similarly apply for splicing patterns, that is, we are interested in total weight of *S* − *T* paths containing a specific splicing pattern as a confidence interval. Since a transcript consisting of *s* exons is a combination of *s* − 1 consecutive junctions, we are interested in splicing patterns that are defined by co-occurrence of *k* disjoint junctions. This motivates the following formal definition of the subgraph quantification problem in graph theory language, generalizing the transcript confidence interval calculation to splicing patterns:

**Definition 1 (AND-Quant)** *Let G* = (*V, E*) *be a directed acyclic graph with an edge flow, and* {*E*_*k*_} *be a list of edge sets with the well-ordering property: If a path visits e*_*i*_ *∈E*_*i*_, *then visits e*_*j*_ *∈ E*_*j*_ *at a later step, i < j. An S* − *T path is “good” if it intersects each E*_*k*_. *For a flow decomposition of G, the total “good flow” is the total flow from good S* − *T paths*. AND-Quant(*G*, {*E*_*k*_}) *asks for the minimum and maximum total good flow for all decompositions of G*.

For quantifying a full transcript, *E*_*k*_ consists of single-edge sets, each corresponding to an edge in the *S* − *T* path representing the transcript. The range of optima for this transcript is exactly AND-Quant(*G*, {*E*_*k*_}). For the aforementioned “local quantification”, the definition of {*E*_*k*_} depends on the specific construction of the graph quantification instance, but in general it is easy to construct one *E*_*i*_ for each junction which satisfies the well-ordering property.

To solve AND-Quant, the first step is to define a similar problem called OR-Quant as follows:

**Definition 2 (OR-Quant)** *Let G* = (*V, E*) *be a directed acyclic graph with an edge flow, and E*′ *be an arbitrary subset of E. An S* − *T path is “good” if it intersects E*′. *The total “good flow” and the objective are defined in the same way as in* AND-Quant

The OR-Quant problem is complementary to AND-Quant and is interesting on its own, as it is suitable to represent analyses where we are interested in aggregated expression that includes any of the several junctions or exons. We now convert an instance of AND-Quant to OR-Quant by constructing a graph *G*_*B*_, called the block graph, that contains every edge that is either in one of *E*_*k*_, or is between an edge in *E*_*k*−1_ and an edge in *E*_*k*_ for 1 *≤ k ≤ m*, where *m* is the number of sets in {*E*_*k*_} (with some abuse of notion, assume *E*_0_ consists of a virtual edge ending at *S*, and *E*_*m*+1_ consists of a virtual edge starting at *T*). We claim the following for AND-Quant:

#### Lemma 1

*An S* − *T path is “good” in the sense of Definition 1 if and only if it is a subgraph of G*_*B*_.

If we know the minimum and maximum total “bad flow” (negative of good flow), we can obtain the answer to AND-Quant by complementing the result with *U*, the total flow of *G*. From the lemma, an *S* − *T* path is bad if and only if it intersects with *G* − *G*_*B*_, which turns the problem into an instance of OR-Quant

We now solve OR-Quant. Recall *E*′ is the set of edges that a “good” path needs to intersect. To that end, we define two auxiliary graphs. *G*^−^ is a copy of *G* without the edges in *E*′. *G*^+^ is a copy of *G* with an extra vertex *T* ′ which replaces *T* as the flow sink, and all edges in *E*′ have their destination moved to *T* ′. We claim running MaxFlow on both graphs yields the solution:

**Theorem 1** *Let G*^+^ *and G*^−^ *be constructed as described above. Then* OR-Quant(*G, E*′) = [*U* − MaxFlow(*G*^−^), MaxFlow(*G*^+^)].

Constructing *G*^+^ and *G*^−^ takes linear time, so the time complexity of OR-Quant depends only on the time to solve the MaxFlow. Combining the arguments leads to the solution to AND-Quant (recall *U* is the total flow on *G*):

**Theorem 2** *Let G*_*B*_ *be the block graph, and* [*l, r*] = OR-Quant(*G, G* − *G*_*B*_). *Then* AND-Quant(*G*, {*E*_*i*_}) = [*U* − *r, U* − *l*].

See Figure 2 for example of both problems. Algorithm 1 shows the pseudocode for both OR-Quant and AND-Quant. The proofs of the above theorems can be found in Section S2–S3.

**Figure 2:**
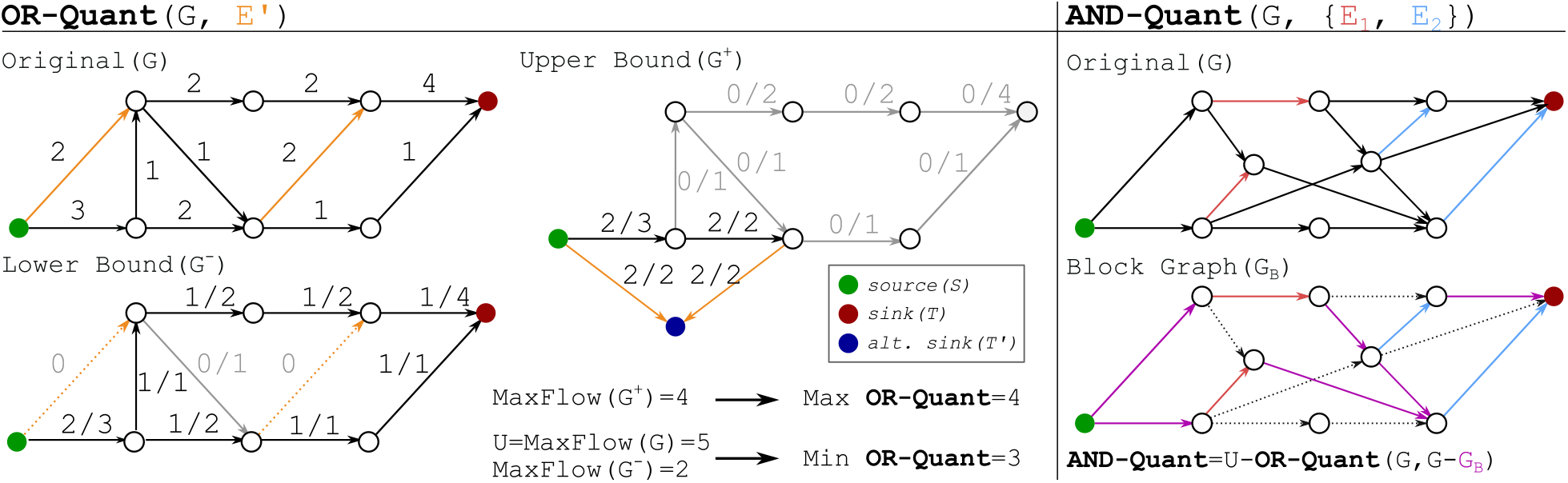
Examples for Subgraph Quantification. Left: Example for OR-Quant, showing *G* and the auxiliary graphs *G*^+^ and *G*^−^. Right: Example for AND-Quant, showing *G* and the block graph *G*_*B*_. As noted in the figure, *G*_*B*_ consists of all colored / non-dashed edges in the image.

### 2.5 Structured Analysis of Differential Expression

We have discussed non-identifiability-aware transcript quantification under two assumptions. In this section, we model the quantification problem under a hybrid assumption: Some fragments are generated from the reference transcriptome while others are generated by combining known junctions (valid under the setup of graph quantification). This model is more realistic than the model under the two extreme assumptions.

For each transcript *T*_*i*_, let 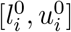 denote its range of optimal expression calculated under the complete reference transcriptome assumption (as in Section 2.2), and 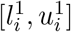 denote its range of optimal expression calculated under incomplete reference transcriptome assumption (as in Section 2.4). We use parameter *λ* to indicate the assumed portion of fragments generated by combining known junctions. For 0 *< λ <* 1, we define 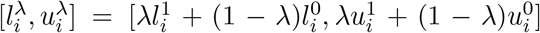, interpolating between the ranges for the extreme assumptions. These ranges are useful for analyzing the effects of non-identifiability under milder assumptions, as we now show for differential expression.

In differential expression analysis, for each transcript we receive two sets of abundance estimates {*A*_*i*_}, {*B*_*i*_} under two conditions, and the aim is to determine whether a transcript is expressed more in {*A*_*i*_}. With fixed *λ*, we can instead calculate the ranges 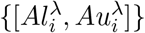 and 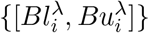 as described above. Suppose the transcript is detected to be differentially expressed by comparing {*A*_*i*_} and {*B*_*i*_} and it is over-expressed in {*A*_*i*_}. When considering non-identifiability, if the mean of 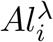 is less than that of 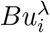 by a threshold, we define this transcript to be a questionable call for differential expression. If *λ* portion of expression is explained by unannotated transcripts, we cannot determine definitely if *A*_*i*_ is larger on average than *B*_*i*_. This filtering of differential expression calls is very conservative (expected to filter out few calls), as most differential expression callers require much higher standards for a differential expression call.

## 3 Results

### 3.1 Implementation

In the following analyses, we use Gencode annotation (version 26) [30] as the reference transcriptome for constructing the splice graphs and for expression quantification. Salmon [4] and its corresponding graph quantification probabilistic model [23] are used to obtain the edge abundances of the splice graphs under complete and incomplete reference assumption. We evaluate the expression estimation uncertainty due to non-identifiability by determining whether the ranking of expression can be altered under different optimal abundances. Specifically, the ranking is computed within the same sample across isoforms in Section 3.2, and for the same isoform across samples in Section 3.3, which is statistically defined by differential expression (DE) analysis.

### 3.2 Expression of 20%–47% Transcripts Have Inconclusive Ranking Compared to Sibling Isoforms

We applied our methods to the Human Body Map dataset [31] (SRA accession ERX011205), which consists of 16 RNA-seq samples from 16 tissues. We are interested in evaluating the expression estimation uncertainty due to non-identifiability, and focus on the transcripts for which the uncertainty of expression estimation is so large that the ranking of expression between the transcript and its sibling isoforms cannot be determined. We use the term “sibling isoforms” of a transcript to refer to the annotated alternative splicing isoforms that belong to the same gene. For each transcript, we enumerate its sibling isoforms, and compare the range of optimal expression estimates for the pair of transcripts to determine whether the two ranges overlap. An overlap between the two ranges indicates an indecisiveness in the ranking of expression between the two transcripts.

We observe that around 35%–50% of transcripts have uncertain expression estimates due to non-identifiability. That is, the ranges of optimal abundances of these transcripts are not single points. The majority of them (around 20%–47% of total transcripts across all 16 samples) have a very uncertain expression estimate such that the ranking of expression between the transcript and at least one of its sibling isoform is inconclusive (Figure 3A). The range is computed under various *λ* values (compositions of reference transcripts expression) ranging from 10% to 100%. These transcripts are possibly false positives in isoform switch detection if they are predicted.

**Figure 3:**
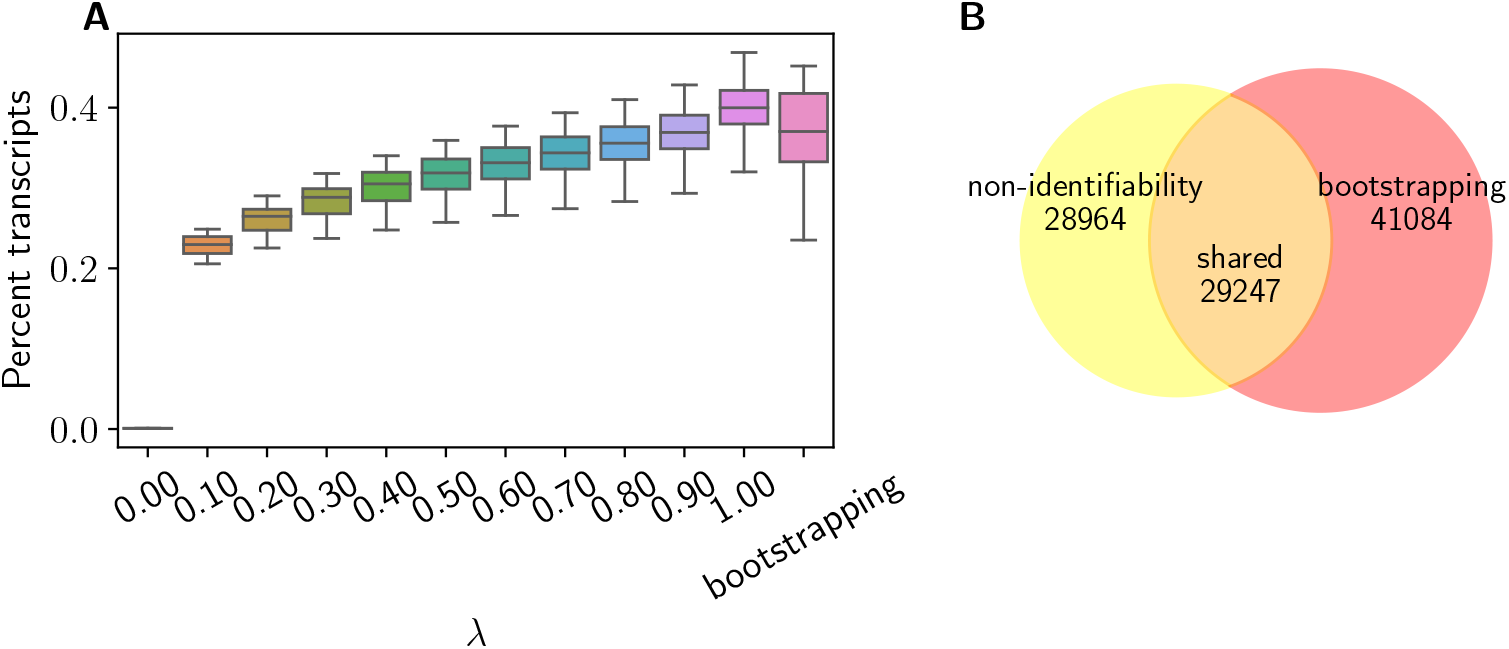
(A) Percentage of transcripts that have inconclusive ranking of expression compared with their sibling isoform due to model non-identifiability under various composition of splice graph paths expression. X-axis indicate the expression composition of splice graph paths. The last column of x-axis represents another source of expression uncertainty, finite sample size, which is evaluated by the bootstrapping method implemented in the Salmon quantifier. (B) Venn diagram showing the unique and overlapping transcripts with inconclusive ranking of expression due to model non-identifiability and due to finite sample size. This subplot corresponds to the prostate sample of Human Body Map dataset. *λ* is set to be 70% for model non-identifiability case.

When compared with the expression estimation uncertainty caused by finite sample size (or finite sequencing depth), we observe that the sets of transcripts with inconclusive expression ranking due to the two sources of error do not have large overlap (Figure 3B). In an arbitrarily chosen Human Body Map sample (a prostate sample, SRA accession ERR030877), we set the *λ* value to be 70% to make the number of transcripts with uncertainty expression estimates under the two cases similar. Only half of transcripts for which the uncertainty estimates are caused by finite sample size are common to the transcripts under the model non-identifiability case. This observation suggests that the expression uncertainty due to model non-identifiability cannot be captured by the degree of uncertainty due to finite sample size. The uncertainty caused by finite sample size is evaluated by bootstrapping in Salmon software [4]. Terminus [32] identifies groups of transcripts that have smaller quantification variance when quantifying as a group compared to quantifying individually, and is based on bootstrapping or Gibbs sampling of expression estimation. Theoretically, the difference between Terminus and our approach for identifying uncertain expression estimates lies in the source of error and whether incomplete reference is considered. Figure S2 shows the percentage of commonly identified transcripts with expression estimation uncertainty of the two methods in Human Body Map dataset.

The computed percentages of transcripts with inconclusive ranking of expression are upper bounds. Because the range of optimal expression estimate per transcript is calculated separately, arbitrarily selecting a pair of values from two ranges of optimal expression of two transcripts may not lead to an optimal pair of expression in the expression probabilistic model. For example, selecting the maximum values for both isoforms may lead to non-valid estimates where the sum of estimated expression (before normalization) exceeds the number of observed RNA-seq reads. However, we speculate that these upper bounds are close to the true percentages. Because reverting the ranking requires one to increase the expression of one transcript and decrease the expression of the other, and it is less likely to generate a non-viable paired estimates under this operation.

### 3.3 Differentially Expressed Transcripts Are Generally Reliable When Assuming the Reference Transcripts Contribute More Than 40% to the Expression

Applying our method to the MCF10 cell line samples with and without EGF treatment (accession SRX1057418) [33], we analyze the reliability of the detected differentially expressed (DE) transcripts. We use Salmon [4], tximport [11], and DESeq2 [9] for the differential expression detection pipeline. This pipeline predicts 257 DE transcripts under a FDR threshold of 0.01. We use the overlap between the mean ranges of optimal expression estimated in the samples with and without EGF treatment as an indicator for unreliable DE detection (see Section 2.5 for details). The overlap is defined as the size of the intersection over the size of the smaller range of the two ranges of optima. We compute the overlap under various *λ* values. The threshold to declare an unreliable DE detection is 25% of overlap.

We identify examples of reliable and unreliable DE predictions. There are five differentially expressed transcripts for which their differential expression statuses may change even when reference expression proportion (1 − *λ*) is as high as 90%. Figure 4A–C shows the lower bounds and upper bounds of the transcript expression of three examples among the five transcripts. Their expression estimates suffer from great uncertainty such that the ranges of optima between the two DE groups overlap. The five genes corresponding to the five transcripts are involved in the following cellular processes or pathways: mRNA degradation, cell apoptosis, glucose transportation, DNA repair, inhibition and transportation of certain kinetics [34]. These genes contain between 6 and 22 isoforms. Further analyses based on the DE detection of these five transcripts require caution since they may be falsely detected to be differentially expressed due to non-identifiability.

**Figure 4:**
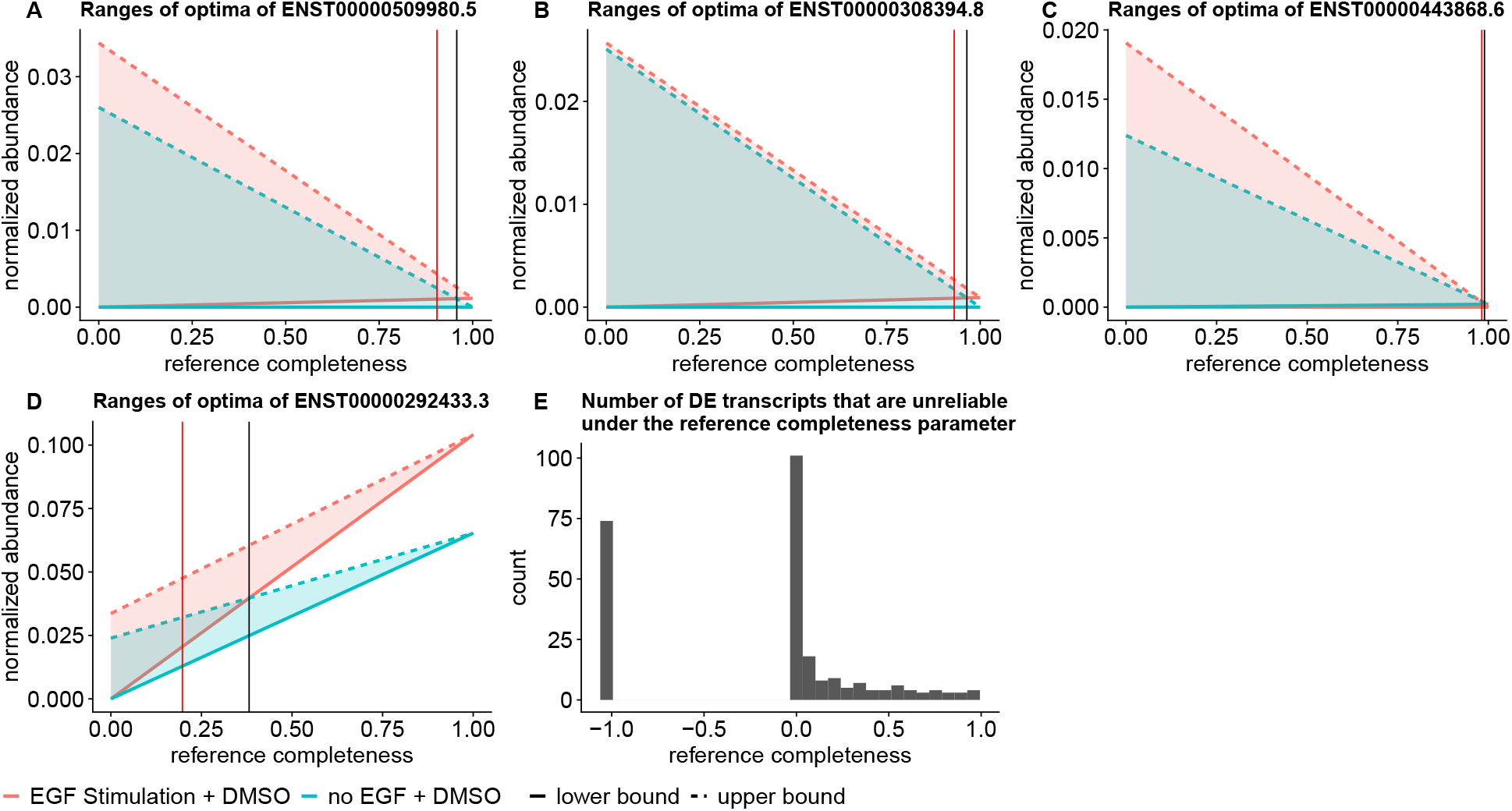
(A–D) Mean ranges of optima of DE groups. X axis is assumed reference completeness 1 − *λ*, the proportion of expression from the reference transcriptome. Y axis is the normalized abundances, where Salmon estimates are normalized into TPM for linear programming under complete reference assumption, and total flow in subgraph quantification is normalized to 10^6^ for each sample. Black vertical lines indicate the reference completeness where the mean ranges of optima overlap. Red vertical lines indicate the reference completeness that the ranges have 25% overlap. (A) The potential unreliable DE transcript ENST00000509980.5. (B) The potential unreliable DE transcript ENST00000308394.8. (C) The reliable DE transcript ENST00000443868.6. (D) The reliable DE transcript ENST00000527470.5. (E) The histogram of the number of unreliable DE transcripts at each *λ* value. Unreliability is defined as more than 25% overlap of the ranges of optima. − 1.0 in the x axis indicates the overlap is no greater than 25% over all *λ* values.

Other than these transcripts, the detected DE transcripts are generally reliable after considering the potential expression estimation uncertainty due to non-identifiability. The ranges of optimal expression estimates between the two sample groups do not have large overlaps when the reference transcriptome is relatively complete and contribute more than 40% to the expression (Figure 4E). This observation is supported by another dataset, where replicates of naive CD8 T cells and four replicates of effector memory CD8 T cells are compared for differential expression detection (Figure S3). There are 3152 differentially expressed transcripts under FDR threshold 0.01, 19 out of which are unreliable even when reference transcripts compose more than 90% expression. We observe the similar pattern that most DE prediction are reliable when the reference transcript expression is more than 40%.

A previous study [35] showed that expression quantification software tend to make slightly more mistakes in deciding the relative expression of isoforms within one sample, compared to deciding the fold change of one isoform across multiple samples. Our results here and in the previous section show agreement with that observation, but with amplified errors in deciding relative expression of isoforms within one sample. Our model for bounding the range of uncertainty due to model non-identifiability provides an explanation from the theoretical perspective: the short sequencing reads may not be sufficient to uniquely reveal the relative abundances of transcripts from complicated splice graphs.

## 4 Conclusion and Discussion

We develop algorithms to compute the range of optimal expression estimates due to non-identifiability for each transcript. We consider both complete and incomplete reference transcripts assumption (quantified with with reference-transcript-based quantification and graph quantification, respectively) for method development, and further provide the range of uncertain estimates under mixed assumptions: a certain proportion of expression is from reference transcripts and the rest (indicated by *λ*) is from expression of splice graph paths. Applying our methods on Human Body Map samples and two RNA-seq datasets for DE transcripts detection, we observe the following expression uncertainty patterns: the ranking of expression between a transcript and its sibling isoforms in a given sample cannot be determined for many (20%–47%) transcripts if the expression estimation uncertainty is considered; but when comparing the expression estimates of a transcript across multiple RNA-seq samples, the detected differentially expressed transcripts are mostly reliable.

The *λ* parameter is unknown in our model, and we address this by investigating the ranges of optima under varying reference completeness values. However, determining the best *λ* value that fits the dataset as an indicator for reference trustfulness is an interesting question in itself, and we believe transcript assembly or related methods might be useful for choosing the correct *λ* value for each dataset.

The non-identifiability problem in expression quantification is partly due to the contrast between the complex splicing structure of the transcriptome and short length of observed fragments in RNA-seq. Recent developments of full-length transcript sequencing might be able to close this complexity gap by providing longer range phasing information. However, full-length transcript sequencing techniques suffer from problems such as low coverage and high error rate. It is still open whether full-length transcript sequencing is appropriate for quantification and how the current expression quantification methods, including this work, should be adapted to work with full-length transcript sequencing data.

## Acknowledgements

This work has been supported in part by the Gordon and Betty Moore Foundations Data-Driven Discovery Initiative through Grant GBMF4554 to C.K., by the US National Institutes of Health (R01GM122935), and the US National Science Foundation (DBI-1937540). This work was partially funded by The Shurl and Kay Curci Foundation. This project is funded, in part, under a grant (#4100070287) with the Pennsylvania Department of Health. The Department specifically disclaims responsibility for any analyses, interpretations or conclusions.

## Disclosure Statement

C.K. is a co-founder of Ocean Genomics, Inc.

## Supplementary Information

### S1 Pseudocode for OR-QUANT and AND-Quant

**Algorithm 1** Algorithms for OR-Quant and AND-Quant

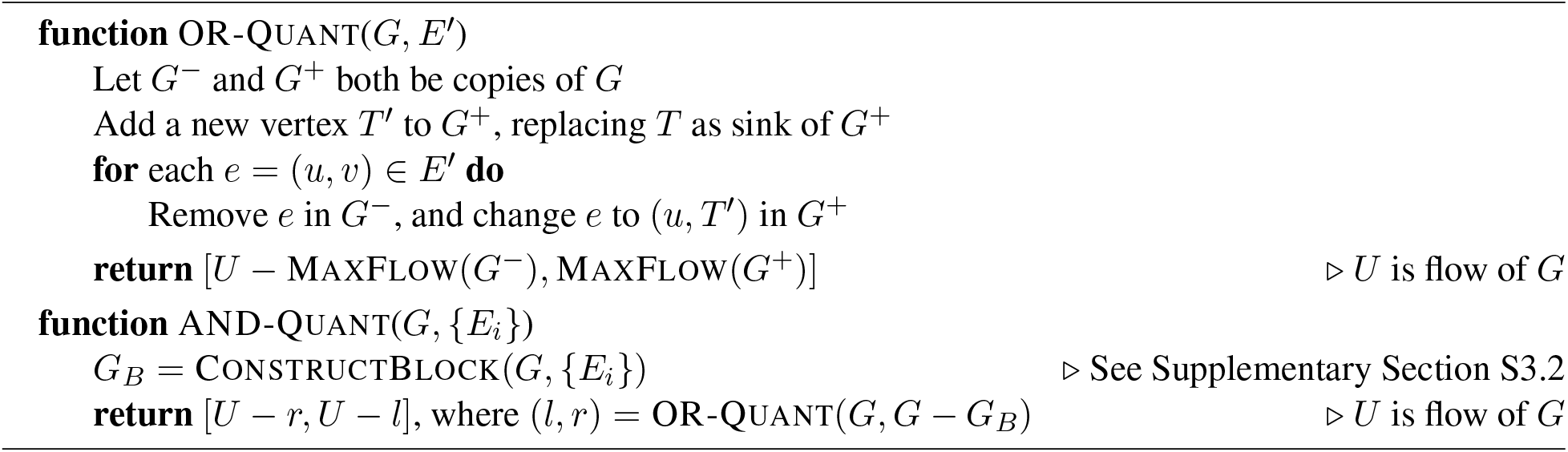

### S2 Algorithms for OR-QUANT

In this section, we state the algorithms for OR-Quant from a pure graph theoretical perspective. Recall our setup consists of a DAG *G* with predetermined source *S* and sink *T*, a flow *F* = {*f*_*e*_} on the graph, and an edge subset. We also use *C*(*F*) to denote the total flow, and in this way we write *U* = *C*(*F*).

We start by several basic definitions.

**Definition S3 (Flow)** *Given DAG G with predetermined source S and sink T, a flow F is a set of edge weights of G satisfying non-negativity on every edge weight and flow balance on every vertex except S and T*.

**Definition S4 (Decompositions)** *Given graph G and a flow*, *F =* {*f*_*e*_} *a decomposition R is written as* {*T*_*i*_, *c*_*i*_}*where each T*_*i*_ *is an S* − *T path on and is a nonnegative number, satisfying* 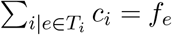 *for every edge e. We use* |*R*| *to denote the number of S* − *T paths in R, and C*(*R*) = Σ *c*_*i*_ *to denote the total flow / capacity of the decomposition*.

**Definition S5 (Partial Decompositions)** *A partial decomposition of a set of non-negative edge weights W* = {*w*_*e*_}*on a graph is written as where each is again an S* − *T path on G, and c*_*i*_ *is a nonnegative number satisfying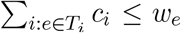 for every edge e. Alternatively, if W is a flow, a partial decomposition can simply be defined as a subset of a (full) decomposition of F*.

Note that we define partial decompositions on arbitrary non-negative edge weights, as required for a later proof. We present the following lemma without proof.

#### Lemma S2

**(Finite Decomposition)** *Every flow on a graph of finite size has a decomposition of finite size*.

We now restate the definition of OR-Quant slightly more formally.

**Definition S6** *Let G* = (*V, E*) *be a DAG with a flow F, and E* ′ ⊆*E. An S* − *T path is good if it intersects E* ′. *(We also say the path is bad if it does not intersect E* ′.*) For a decomposition, the total good flow is defined as* 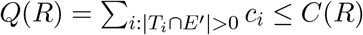, *that is total flow from good paths in the decomposition*. OR-Quant(*G, E* ′) *asks for the minimum and maximum Q*(*R*) *for any possible decomposition R of F*.

We now state our algorithms and prove the correctness of the algorithms, starting from the lower bound.

**Theorem S3 (Diff-Flow)** *The auxiliary graph G*^−^ *is built with edge set E* −*E* ′. *Z* = *U* MaxFlow(*G*^−^) *is the minimum total good flow Q*(*R*) *from any possible decomposition R of F*.

**Proof:** We prove the theorem in two parts: First, we show there is a decomposition *r* such that *Q*(*R*) = *Z*, then we show any decomposition satisfies *Q*(*R*) *≥ Z*.

The key observation is that all *S* − *T* paths that are bad are fully in *G*^−^. For the first part, fix a maximum flow of *G*^−^ as *F* ^−^. We let *R*^−^ be an arbitrary decomposition of *F* ^−^, and *R*^+^ be an arbitrary decomposition of *F* − *F* ^−^.

We have *Q*(*R*^−^) = 0 because no path in *R*^−^ intersects with *E*′. We now claim *Q*(*R*^+^) = *Z*. Because *C*(*R*^+^) = *C*(*R*) − *C*(*R*^−^) = *Z, Q*(*R*^+^) = *Z* if and only if every path in *R*^+^ intersects with *E*′. Assuming otherwise, we can remove that path from *R*^+^ and add it to *R*^−^, which leads to larger *C*(*R*^−^). This is impossible: it implies *F* ^−^ is not the MaxFlow of *G*^−^, because the flow of *R*^−^ is a strictly better solution. Having proved *Q*(*R*^+^) = *Z*, we now know *R*^−^ + *R*^+^, a (full) decomposition, has exactly *Z* total good flow.

Now for any decomposition *R*, we split it into two parts. *R*^+^ contains all good paths in *R*, and *R*^−^ contains all bad paths in *R*. We have *C*(*R*^−^) *≤* MaxFlow(*G*^−^), because the flow of *R*^−^ is a flow on *G*^−^. Because *Q*(*R*^−^) = 0, we have *Q*(*R*) = *Q*(*R*^+^) = *C*(*R*^+^) *≥ Z*.

**Theorem S4 (Split-Flow)** *The auxiliary graph G*^*^ *is built by adding a vertex T* ′ *in G and, for each edge in E* ′ *changing its destination to T* ′ *(this was called G*^+^ *in main text, but we change it to G*^*^ *for technical reasons here). T* ′ *replaces T as the sink of flows. Y* = MaxFlow(*G*^*^) *is maximum total good flow for any possible decomposition of F*.

**Proof:** We prove the theorem in a similar style: First we show there is a decomposition *R* of *F* such that *Q*(*R*) = *Y*, then we show any decomposition satisfies *Q*(*R*) *≤ Y*.

For the first part, we first build an arbitrary decomposition *R*^*^ from a maximum flow of *G*^*^. However, *R*^*^ is not a valid partial decomposition of *F*, because *G*^*^ and *G* have different sets of edges. Our goal is to obtain a partial decomposition *R*^+^ of *F* that is a “reconstruction” of *R*^*^. We will define several terms for convenience of our proof.

#### Path Mapping for *G*^*^

A good *S* −*T* path on *G* is mapped to an *S T* ′ path on *G*^*^ by finding the first edge of the path that is in *E*′, change the destination of that edge to *T* ′, and discard the edges after that so the path ends at *T* ′. (We will never map a bad *S* − *T* path on *G* this way.)

Similarly, an *S* − *T* ′ path on *G*^*^ is mapped to a (incomplete) path on *G* by moving the destination of last edge (that was *T* ′) back to its original node before the transformation. We assume a label containing its original destination is kept on each edge that ends at *T* ′. This implies multiple edges can exist between a node and *T* ′, and the movement follows the label. The resulting path is guaranteed to intersect with *E*′, but is not a complete *S* − *T* path.

We apply an induction argument, formally defined as follows:

#### Flow Reconstruction Instance

An instance for flow reconstruction has two inputs: (*F, R*^*^). *F* is a flow on *G*, and *R*^*^ is a partial decomposition on *F* ^*^ (of *G*^*^), where *F* ^*^ is constructed in the same way as *G*^*^ by moving endpoints of edges in *E*′ to a new node *T* ′. *F* ^*^ is actually not a flow because load balance is violated at the vertices that has an incoming edge moved away. Nevertheless, partial decomposition of *F* ^*^ is still well-defined as *R*^*^ contains *S* − *T* ′ paths. The output of the instance is *R*^+^, a partial decomposition of *F*, satisfying that (1) Each path in *R*^+^ is good and (2) If we map each path in *R*^+^ back to *G*^*^, the resulting partial decomposition has the same flow as *R*^*^.

We will apply the induction on |*R*^*^|, number of paths in *R*^*^. The base case is then |*R*^*^| = 0, where we can simply return an empty *R*^+^. Assume now the flow reconstruction can be solved for |*R*^*^| *≤ k* − 1, we solve the instance with |*R*^*^| = *k*. We first state the algorithm, then prove its correctness.

#### Induction Algorithm

We first pick the path (*P, c*) in *R*^*^ whose second-to-last vertex (right before *T* ′) is the last among all paths in topological ordering of *G*. Denote the last edge of this path, mapped back to *G*, as (*u, v*). *P* without last edge is then a path from *S* to *u*. We let *P* ′ denote *P* with last edge changed back to (*u, v*).

We next run a maxflow on *G* from *v* to *T* with total flow *c*. This call always return a full flow (with total flow *c*), because there are at least *c* incoming flow for *v* from the existence of edge (*u, v*) alone. This flow is then decomposed, and for each path in the decomposition, we prepend it with *P* ′ (which is a path from *S* to *v*) so it becomes a *S − T* path. We call the resulting partial decomposition *R*^0^, and we next show it is valid, meaning the total flow of *R*^0^ on each edge does not exceed *F* . For edges in *P* ′, the total flow on this edge in *R*^0^ is exactly *c*. As (*P, c*) is a path in *R*^*^, each of the edges in *P* has at least *c* flow in *F* ^*^, which immediately means each edge in *P* ′ has at least *c* flow in *F* . For all other edges, its flow comes from the *v* − *T* maxflow, so the flow value is always below the capacity of this edge.

We continue the process by recursively calling the procedure on (*F* ′, *R*′^*^), where *F* ′ is *F* minus the flow of *R*, and *R*′^*^ is *R*^*^ without (*P, c*). We return the partial decomposition from the recursive call merged with *R*^0^.

#### Proof of Induction Correctness

The new instance to be called is an instance with size *k* − 1, since one path in *R*^*^ is removed. We first prove that the recursive call is valid, that is, *R*′^*^ is indeed a partial decomposition of *F* ′^*^ (*F* ′^*^ is constructed from *F* ′ by moving the endpoint of edges in *E*′ to *T* ′).

If the call is invalid, it means some edge in *F* ′^*^ has less flow than total flow from *R*′^*^. Since *R*^*^ is a partial decomposition of *F* ^*^, the condition will only be invalidated for an edge with positive flow in *R*^0^, as these are the only edges where the flow on *F* ′^*^ is less than the flow on *F* ^*^. This edge can either be an edge in *P*, or an edge whose starting point is later than *v* in the topological order of *G* by construction of *R*^0^ (with a *v* − *T* maxflow). If the edge is in *P* and is not the last edge, exactly *c* flow is subtracted on this edge from *F* to *F* ′ and from *R*^*^ to *R* ′^*^, so this would contradict with the condition that *R*^*^ is a partial decomposition of *F* ^*^. If this is the edge (*u, T* ′) that is mapped from (*u, v*), again exactly *c* flow is subtracted in both *F* to *F* ′ and *R*^*^ to *R*′^*^.

If this is an edge not in *P*, we claim the edge has zero flow in *R*′^*^. This is because the way we pick (*P, c*), where the path with latest second-to-last vertex is picked. If the edge has positive flow in *R*′^*^, it implies there is a path in *R*′^*^ that includes this edge, and the ending point of the path would be later than *v*, meaning the path would have been chosen in place of *P* during the process.

Given the recursive call is valid, we can prove the correctness of the algorithm by showing that *R*^+^ outputted by the algorithm indeed satisfies the two output conditions. For (1), each path in *R*^+^ is good because it come from a *R*^0^ at some iteration, and all paths in *R*^0^ has *P* ′ as prefix whose last edge (*u, v*) is in *E*′. For (2), at each recursion call, *R*^0^ mapped to *F* ^*^ becomes one path *P* with flow *c*, which matches (*P, c*) in *R*^*^, so the condition is maintained similarly by an induction argument. This finishes the correctness proof for the induction.

#### Completing the Proof

With the induction algorithm, we can reconstruct a partial decomposition *R*^+^ of *G* from *R*^*^ (a decomposition of *F* ^*^), then similar to last proof, construct another partial decomposition *R*^−^ from *F* minus the flow of *R*^+^. By construction of *R*^+^, we have *Q*(*R*^+^) = MaxFlow(*G*^*^). If some path in *R*^−^ intersect with *E*′, we can remove that path from *R*^−^ and add it to *R*^+^, then map the new *R*^+^ to *G*^*^ to obtain a flow better than MaxFlow(*G*^*^), a contradiction. This means *Q*(*R*^−^) = 0 and the decomposition *R* = *R*^+^ + *R*^−^ satisfies *Q*(*R*) = MaxFlow(*G*^*^).

Figure S1 provides a concrete example for the induction argument. For the second part of the statement, for a decomposition *R* of *F*, we can write *R* = *R*^+^ + *R*^−^ where *R*^+^ contains all good paths in *R* and *R*^−^ contains all bad paths in *R*. We can map *R*^+^ to *G*^*^ and denote the resulting partial decomposition *R*^*^ (*R*^*^ is indeed a partial decomposition by noting there is a one-to-one mapping between edges of *F* ^*^ and edges of *F*). We have *Q*(*R*) = *Q*(*R*^+^) = *C*(*R*^+^) = *C*(*R*^*^) *≤* MaxFlow(*G*^*^). This completes the whole proof. □

With both bounds proved, we can formally finish the proof of the main theorem:

**Theorem 1** *Let G*^+^ *and G*^−^ *be constructed as described above. Then*

OR-Quant(*G, E*′) = [*U* − MaxFlow(*G*^−^), MaxFlow(*G*^+^)].

**Proof:** This is implied by Theorem S3 for the lower bound, and Theorem S4 for the upper bound. □

### S3 Algorithms for AND-QUANT

In this section, we prove the complementarity argument mentioned in the main text is correct. Our setup consists of a DAG *G* with predetermined source *S* and sink *T*, a flow *F* = {*f*_*e*_} on *G*, and a list of edge sets {*E*_*k*_} . We define flows and decompositions in the same way as last section (Definition S3, Definition S4). Now recall the definition of AND-Quant, written slightly differently with the new notations:

**Definition S7 (Well-Ordering Property)** *A list of edge sets* {*E*_*k*_} *satisfies the well-ordering property if a path visiting e*_*i*_ *∈ E*_*i*_ *then e*_*j*_ *∈ E*_*j*_ *at a later step implies i < j*.

**Definition S8 (AND-Quant)** *Let G* = (*V, E*) *be a directed acyclic graph with an edge flow, and* {*E*_*k*_} *be a list of edge sets satisfying the well-ordering property. An S* −*T path is good if it intersects each E*_*k*_. *For a decomposition of F, the total good flow is the total flow from good S* −*T paths*. AND-Quant(*G*, {*E*_*k*_}) *asks for the minimum and maximum total good flow for all possible decompositions of F*.

We start by discarding edges *e* in any of *E*_*i*_ such that no good *S* −*T* paths would use *e*. By definition, removal of these edges will not change the answer to AND-Quant as no good *S* −*T* paths would be excluded by removing these edges. This also means each edge *e* in any *E*_*i*_ satisfies that there is a good *S* − *T* path that includes *e*.

Now given *G* is a DAG and the edge sets satisfy the well-ordering property, we can define the following:

**Definition S9 (Start/End-Sets and Natural Order)** *Define U*_*i*_ = {*u*: (*u, v*) *∈ E*_*i*_}, *V*_*i*_ = {*v*: (*u, v*) *∈ E*_*i*_} *and for convenience, V*_0_ = {*S*}, *U*_*m*+1_ = {*T*}.

*We define the natural order u ≤ v if there is a directed path starting at u and ending at v. We define x → V*_*i*_ *as the condition that there is some v*_*i*_ *∈V*_*i*_ *such that x ≤ v*_*i*_ *(intuitively, x can reach V*_*i*_*). Same goes for U*_*i*_ *and the other direction*.

**Definition S10 (The Block Graph)** *Define B*_*i*_, *the i*^*th*^ *block subgraph, the set of edges* (*u, v*) *that satisfies V*_*j*−1_ *→ u and v → U*_*j*_, *for* 1 *≤ i ≤ m* + 1. *The full block graph G*_*B*_ *is the union of all B*_*i*_ *and E*_*i*_.

The filtering steps and construction of the block graph can be done in linear time, as discussed in Section S3.1.

#### Lemma S3

**(Well-Ordering Property Extended)** *If u ∈ U*_*i*_, *v ∈ V*_*j*_, *v ≤ u, then i > j*.

**Proof:** We can derive this from the well-ordering property of {*E*_*k*_}, by constructing an *S* − *T* path. First, as *v ∈ V*_*j*_ there is an edge in *E*_*j*_ that contains *v*, and by our previous assumption there is a path from *S* to *v* that uses this edge. As *v ≤u*, there is a path from *v* to *u*. As *u ∈ U*_*i*_, there is an edge in *E*_*i*_ that contains *u*, and similarly there is a path from *u* to *T* that uses this edge. Combining the three segments together, we have an *S* − *T* path that first visits an edge in *E*_*j*_, then an edge in *E*_*i*_ later, so by the well-ordering property of {*E*_*k*_}, *i > j*. □

#### Lemma S4

*For each v*_*i*_ *∈ V*_*i*_, *v*_*i*_ *→ U*_*i*+1_.

**Proof:** Recall that we removed all edges in *E*_*i*_ that does not belong to any good paths. *v*_*i*_ *∈ V*_*i*_ implies there is an edge (*u*_*i*_, *v*_*i*_) *∈ E*_*i*_ that is contained in a good path, and later in this good path there is an edge in *E*_*i*+1_ that starts at some vertices in *U*_*i*+1_. This implies *v*_*i*_ *→ U*_*i*+1_. □

#### Lemma S5

*For each u*_*i*_ *∈ U*_*i*_, *u*_*i*_ *→ U*_*j*_ *where j ≥ i*. □

**Proof:** *u*_*i*_ *∈ U*_*i*_ means there is an edge (*u*_*i*_, *v*_*i*_) *∈ E*_*i*_, and *u*_*i*_ *→ U*_*i*_. As shown last lemma, *v*_*i*_ *→ U*_*j*_ for *j > i*. This means the same also holds for *u*_*i*_ as *u*_*i*_ *< v*_*i*_. □

We can prove the symmetric statements about *V*_*i*_.

#### Lemma S6

*For a fixed i, each vertex x in G*_*B*_ *either satisfies x → U*_*i*_ *or V*_*i*_ *→ x, but not both*.

**Proof:** If *x* is in one of *U*_*j*_, *x → U*_*i*_ if *i ≤ j* by Lemma S5. Similarly, if *x ∈ U*_*j*_, *V*_*i*_ *→ x* if *i > j* by both Lemma S4.

We can similarly prove that the condition holds for all *x ∈ V*_*j*_, where *x → U*_*i*_ if *j < i*, and *V*_*i*_ *→ x* if *j ≥ i*.

If *x* is in none of *U*_*j*_ or *V*_*j*_, it is in an edge in *B*_*j*_, which by definition of *B*_*j*_ means both *V*_*j*−1_ *→ x* and *x → U*_*j*_. Combined with Lemma S5, this means *x → U*_*i*_ if *j ≤ i*, and *V*_*i*_ *→ x* if *j > i*.

We have proved for any vertex in *x* and any *i*, one of *x → U*_*i*_ or *V*_*i*_ *→ x* must be satisfied. No vertices can satisfy both, otherwise we have *u ∈ U*_*i*_, *v ∈ V*_*i*_ and *v ≤ x ≤ u*, which would imply *i < i* by Lemma S3, contradiction. □

#### Lemma S7

**(Main Lemma for AND-Q****uant**) *An S* − *T path in G is good if and only if it is a subset of G*_*B*_.

**Proof:** We first prove that if a path is good, it is within *G*_*B*_. An edge (*u, v*) on the path is either in some *E*_*i*_, or between some edge in *E*_*i*−1_ and some edge in *E*_*i*_. In latter case, we know *V*_*i*−1_ *→ u* and *v → U*_*i*_, which directly implies (*u, v*) *∈ B*_*i*_.

To prove the other direction, we show that removing any of *E*_*i*_ results in disconnection of *S* to *T* . By the previous lemma, the vertices of *G*_*B*_ can be partitioned into two disjoint sets: 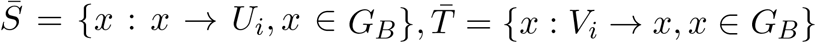. For an edge (*u, v*) *∈ G*_*B*_ that starts in 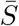 and ends in 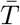:

- If (*u, v*) *∈ B*_*j*_, we know *V*_*j*−1_ *→ u → U*_*i*_ (first part from definition of *B*_*j*_ and second part from 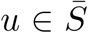, similar for the rest of this proof), so *j* − 1 *< i*, and at the same time *V*_*i*_ *→ v → U*_*j*_ so *i < j*, but there is not an integer *i* between *j* − 1 and *j*, contradiction.
- If (*u, v*) *∈ E*_*j*_, we know *U*_*j*_ *→ u → U*_*i*_ so *j ≤ i* (because *V*_*j*−1_ *→ u* by using the fact there is a good path including *u*), and *V*_*i*_ *→ v V*_*j*_ *→* so *i ≤ j* (because *v ≤ v*_*j*_ *→ U*_*j*+1_ for some *v*_*j*_ *∈ V*_*j*_ by Lemma S4), which implies *i* = *j*.

So we have (*u, v*) *∈ E*_*i*_. In other words, removing all edge in *E*_*i*_ results in disconnection between 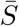 and 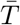, so each *S* − *T* path in *G*_*B*_ uses an edge in *E*_*i*_, for each *i*. This means the path is good. □

We can now finish the proof of the main theorem:

**Proof:** By Lemma S7, a path is a bad path if and only if the path intersects with *G* − *G*_*B*_. Thus, we can convert an instance of AND-Quant to an instance of OR-Quant by constructing the block graph *G*_*B*_, then complementing the results of OR-Quant with *E* ′ = *G* − *G*_*B*_. The correctness of our OR-Quant algorithms is guaranteed by Theorem 1. □

#### S3.1 Constructing *G*_*B*_

In this section we disuss how to construct *G*_*B*_ in linear time. The pseudocode for this algorithm is provided in Section S3.2, and we will be using notations from Section S3.

We first discuss the filtering algorithm. We can first run two breadth-first searches from *S* and *T* to mark the set of vertices that are reachable from *S* and can reach *T* . All vertices that are unable to do both will never appear in an *S* − *T* path, and will be removed first. We let *s*(*v*) denote the largest *i* such that there is a path from *S*, via an edge in each of *E*_1_, *… E*_2_,, *E*_*i*_ in order, to *v*. We now generate the topological order of *G*, and let *s*(*S*) = 0. For every vertex in the topological order other than *S*, the value of *s*(*v*) is determined as follows. *s*(*v*) is initialized as the largest of *s*(*u*) from all its predecessors. Then for every (*u, v*) *∈ E*_*j*_ and *s*(*u*) = *j* − 1, we set *s*(*v*) = max(*s*(*v*), *j*).

We now show the values of *s*(*v*) are correctly derived. If *v* is indeed reachable from *S* via an edge in each of *E*_1_, *… E*_2_,, *E*_*i*_, we have *s*(*v*) *≥ i* as we will visit all nodes on the path in topological order and after an edge in *E*_*i*_ we always have *s*(*v*) *≥ i*. If *s*(*v*) *≥ i*, we show *v* is reachable from *S* via an edge in each of *E*_1_, *E*_2_,, *E*_*i*_. We let *p*(*v*) be the predecessor of *v* where *s*(*v*) is calculated from, either by passing the value or passing an edge in *E*_*j*_. By chaining *p*(*v*), we obtain a path from *S* to *v* where each vertex is responsible for the value of *s*(*v*) of its successor in the path. This means along the path the value will only increase by passing through an edge in *E*_*j*_, and this can only bring *s*(*v*) from *j* − 1 to *j*, so the path must contain an edge from each *E*_1_, *E*_2_, *…, E*_*i*_ in order.

Similarly, we can obtain *t*(*v*) which is defined as the smallest *i* such that there is a path from *v*, via an edge in each of *E*_*i*_, *E*_*i*+1_, *…, E*_*m*_} to *T* by setting *t*(*T*) = *m*+1 and traverse the graph in reverse topological order. The filtering process then works as follows. For each (*u, v*) *∈ E*_*i*_, the edge is kept in *E*_*i*_ if and only if *s*(*u*) = *≥ i* − 1 and *t*(*v*) = *≤ i* + 1. We prove this procedure is correct. A good *S* − *T* path containing (*u, v*) must visit an edge in each of *E*_1_, *E*_2_,, *E*_*i*−1_ in order, visit (*u, v*) *∈ E*_*i*_, then an edge in each of {*E*_*i*+1_, *E*_*i*+2_,, *… E*_*m*_ . The first part is possible if and only if *s*(*u*) *≥ i* − 1, and the last part is possible if and only if *t*(*v*) *i* + 1. It remains to prove that *s*(*u*) *i* 1 (the other part is symmetric). This is because otherwise there will be an edge in *E*_*i*_ that precedes *u*. As (*u, v*) is also in *E*_*i*_, this means there will exist a path containing two edges in *E*_*i*_, which violates the well-ordering property.

We next discuss the block generation algorithm. In fact, we can simply reuse the value of *s*(*v*) and *t*(*v*). For an edge (*u, v*) not in any of *E*_*i*_, if *s*(*u*) + 1 = *t*(*v*), the edge is allocated to *B*_*t*(*v*)_. We prove this procedure is correct. If an edge satisfies *s*(*u*) + 1 = *t*(*v*) = *k*, there is an edge in *E*_*k*−1_ that leads to *u*, meaning *V*_*k*−1_ *→ u*, and similarly *u → U*_*k*_ which is the definition of *B*_*k*_. Similarly, if (*u, v*) *∈ B*_*k*_, we show *s*(*u*) + 1 = *t*(*v*) = *k*. Given *V*_*k*−1_ *→ u* and the edge set is filtered, there exists *v*_*k*−1_ *∈ V*_*k*−1_, *v*_*k*−1_ *≤ v*, and *s*(*v*) *≥ s*(*v*_*k*−1_) = *k* − 1. If *s*(*v*) ≥ *k*, this means there is an edge in *E*_*k*_ that leads to *u* and *V*_*k*_ *→u*. Given *v → U*_*k*_ at the same time, we have *V*_*k*_ → *u ≤ v → U*_*k*_, and there is a path that visits an edge in *E*_*k*_ twice, violating the well-ordering property. We can similarly prove it is necessary and sufficient that *t*(*v*) = *k*.

Combining the results, we have the following lemma for preprocessing (note that topological sorting is a linear time algorithm):

##### Lemma S8

*There is an O*(*n* + *m*) *algorithm that filters* {*E*_*i*_} *such that every remaining edge in every E*_*i*_ *is included in at least one good S* − *T path, and constructs* {*B*_*i*_} *as described in Definition S10, where n and m are the number of nodes and edges in G*.

#### S3.2 Pseudocode for Generating *G*_*B*_

A core component of the AND-Quant algorithm is to compute the block graph *G*_*B*_, which contains all edges that is in some valid *S* − *T* path (that is, a path that visited one edge in *E*_*i*_ in increasing order of *i*). As analyzed in Section S3.1, this can be done efficiently (in linear time), with the pseudocode provided below.

**Algorithm 2** Algorithm to compute the block graph

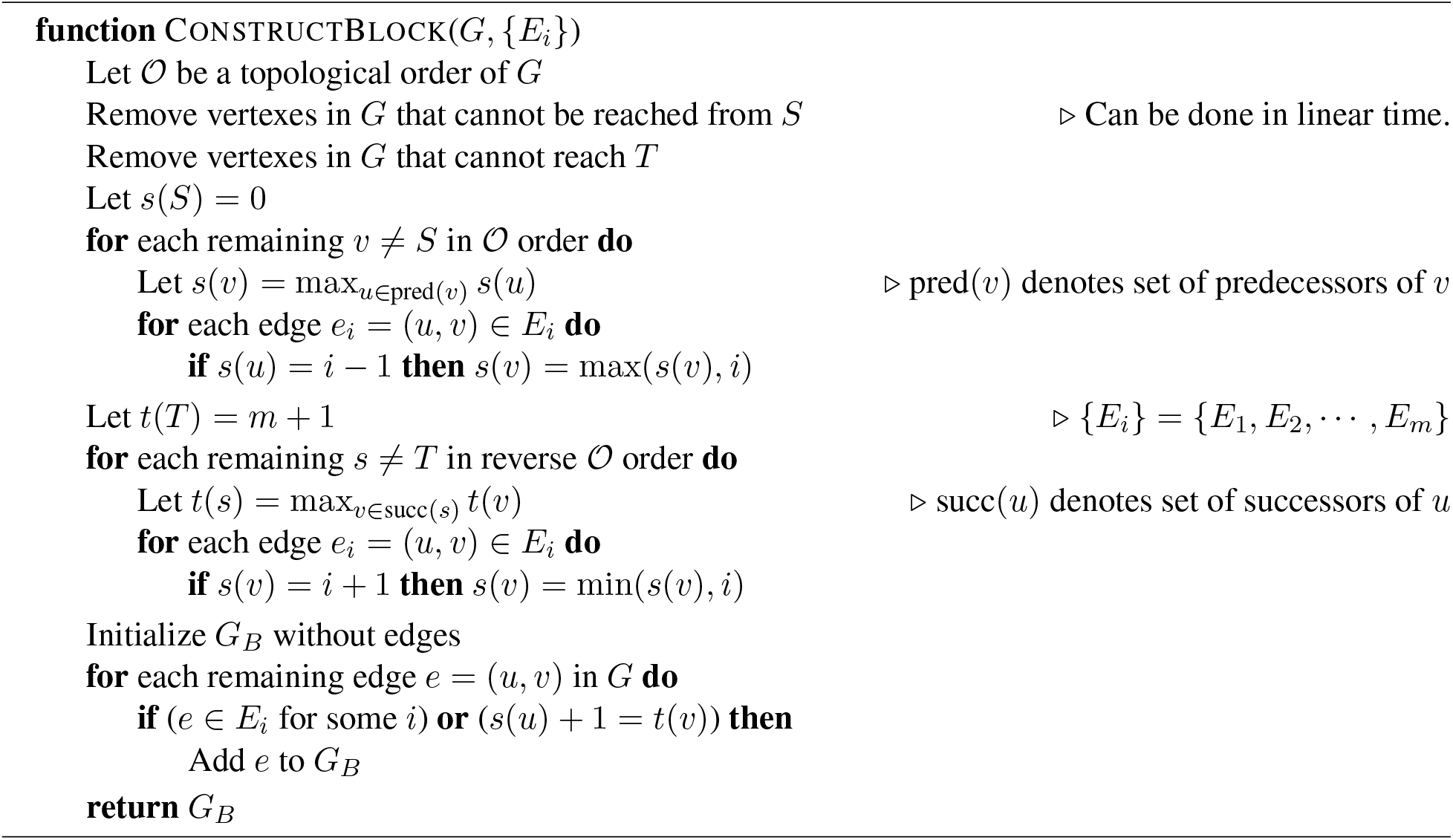

### S4 Supplementary Figures

**Figure S1:**
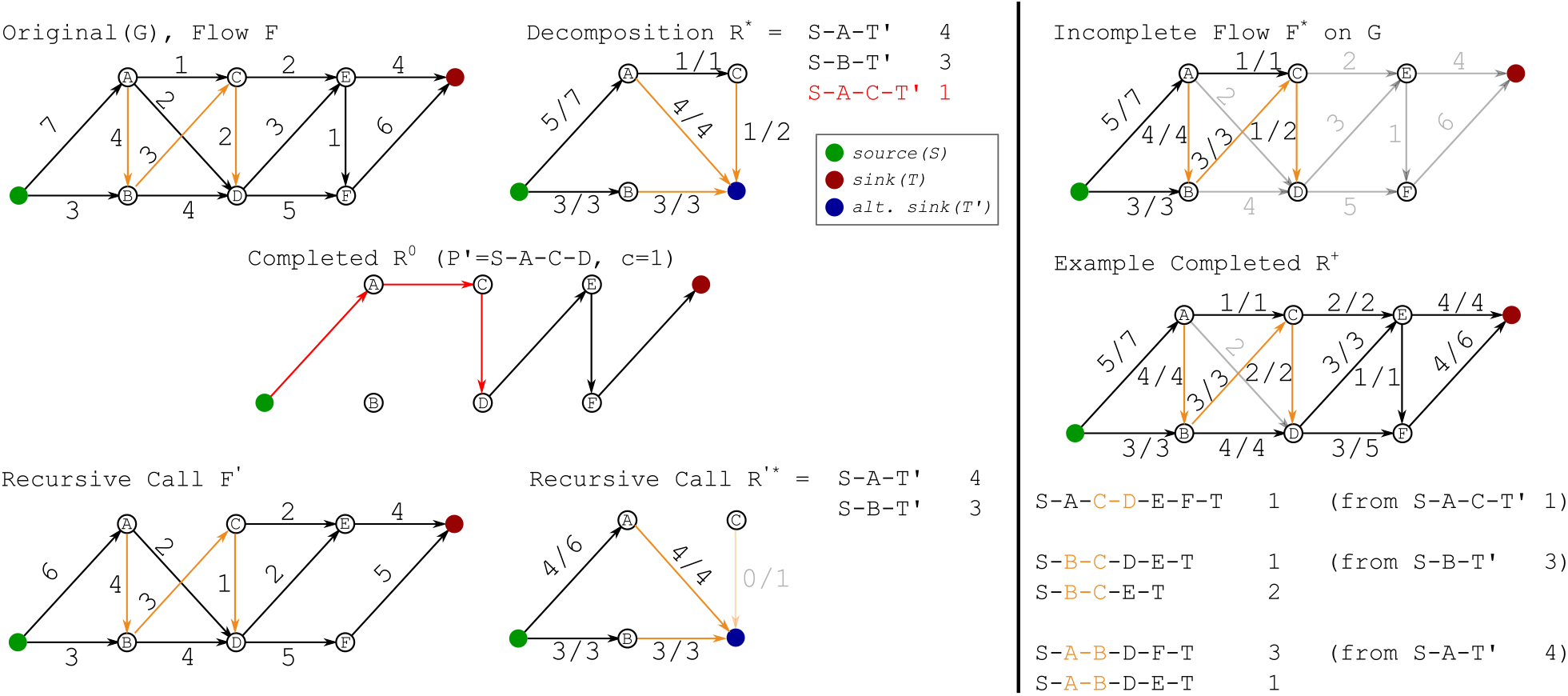
Example for Flow Reconstruction. Input graph *G* is shown on top left, with edges in *E*′ marked in orange. The vertexes *A F* are in topological order. On the **left-hand side** we show one step of the recursive proof. Left top: Input pairs (*F, R*^*^), with the picked path (*P, c*) marked in red. We hide vertices *D, E, F, T* and associated edges in *R*^*^, as those edges cannot be used in *R*^*^. Left middle: Constructed *R*^0^ which contains only one path with weight 1. The part belonging to *P* ′ marked in red. Left bottom: The recursive call (*F* ′, *R*′^*^), which is *F* with *R*^0^ removed and *R*^*^ with (*P, c*) removed respectively. On the **right-hand side** we show a solution to the OR-Quant upper bound. Right top: The maxflow on *G*^*^ directly mapped as *F* ^*^ onto *G* without any completion. Right middle: A completed flow with matching flow value of 8. Right bottom: The solution *R*^+^ obtained from the recursive process. For each path the first edge to appear in *E*′ is marked in orange.

**Figure S2:**
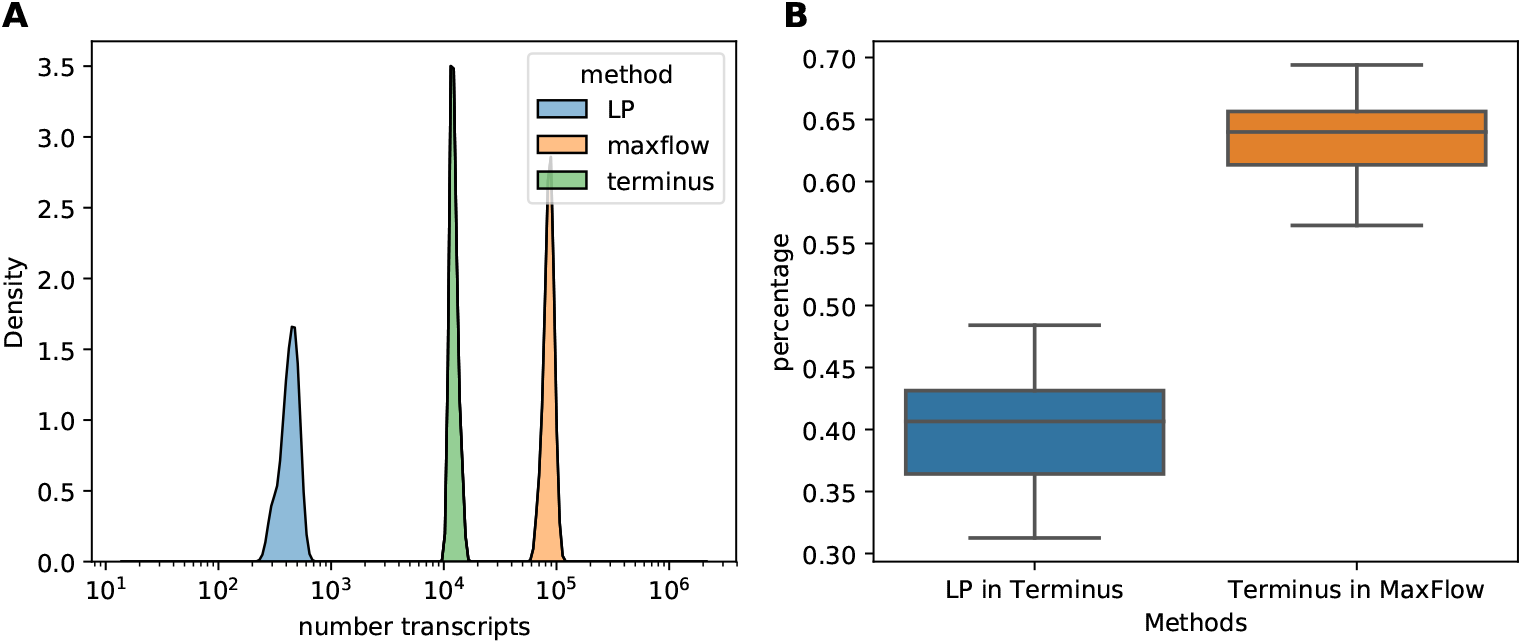
(A) Histogram of number of transcripts identified with uncertain expression estimates under LP, MaxFlowin our approach and under Terminus. (B) Percentage of transcripts identified under LP that are also identified by Terminus, and percentage of transcripts identified by Terminus that are also identified under MaxFlow.

**Figure S3:**
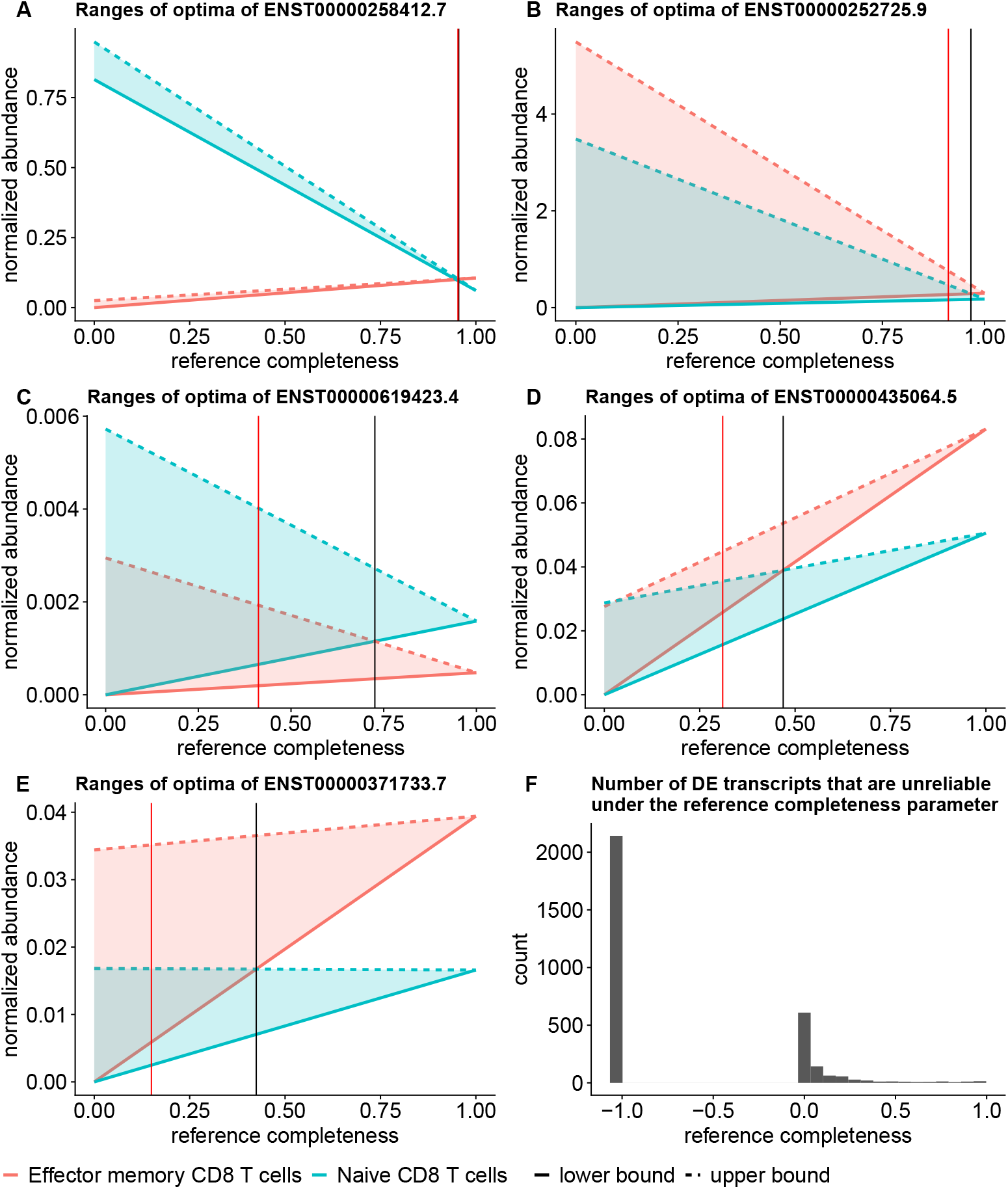
(A–E) Mean ranges of optimal abundances of DE groups. X axis is 1 − *λ*, which is the proportion of expression from the reference transcriptome. Y axis is the normalized abundances, where Salmon estimates are normalized into TPM for linear programming under complete reference assumption, and total flow in subgraph quantification is normalized to 10^6^ for each sample. Black vertical lines indicate the reference completeness where the mean ranges of optima overlap. Red vertical lines indicate the reference completeness that the ranges have 25% overlap. (F) The histogram of the number of unreliable DE transcripts at each *λ* value. Unreliability is defined as more than 25% overlap of the ranges of optima. − 1.0 in the x axis indicates the overlap is no greater than 25% over all *λ* values.

